## Supplementary material for "Reconstructing prehistoric viral genomes from Neanderthal sequencing data": Supplementary Figure S10 Detail of changes and C-T deamination.pdf

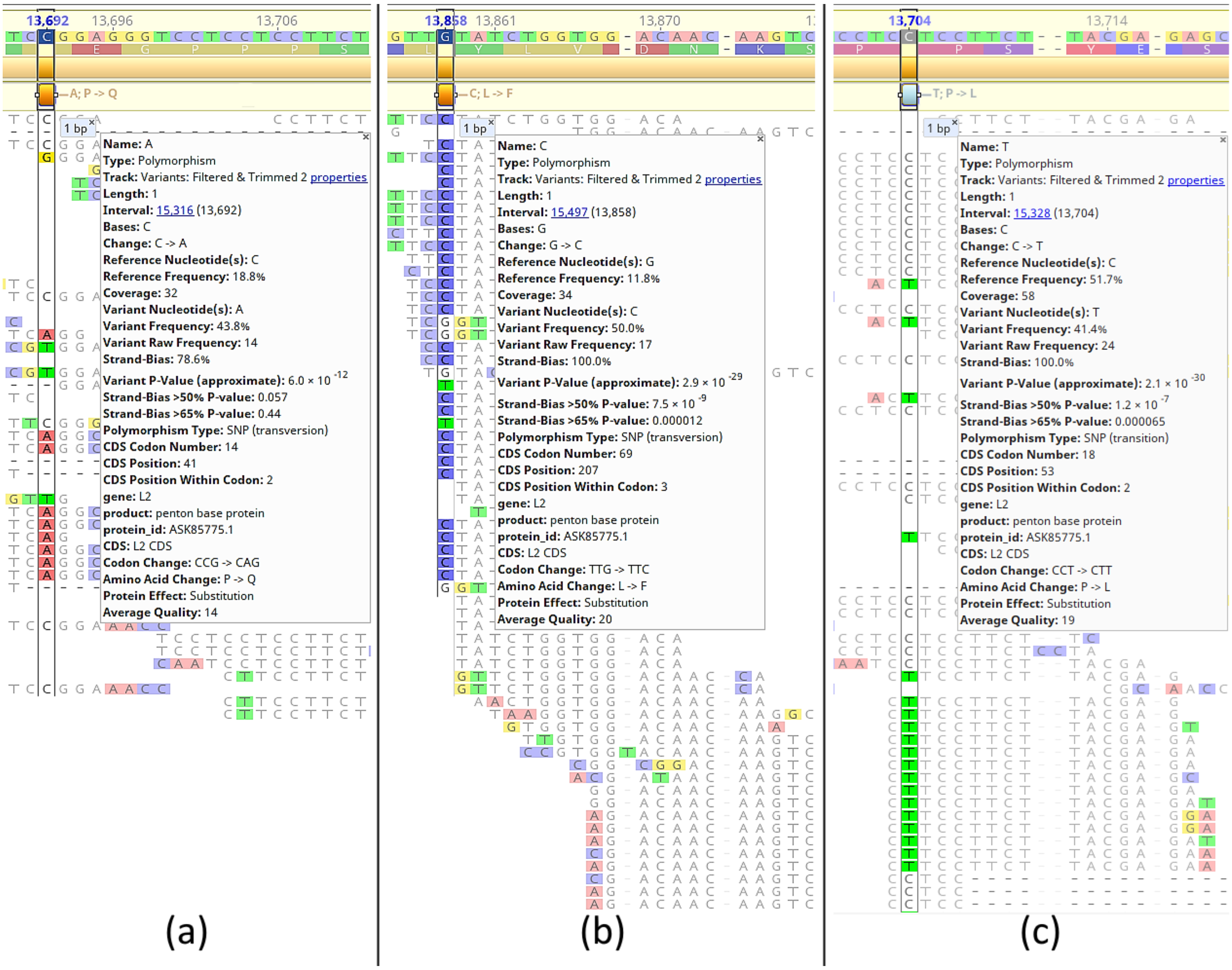


**Supplementary Figure S10.** Details of non-synonymous changes in N-adenovirus sequences (gene L2 - Penton base) with the corresponding statistics and C-to-T changes. In (a) A change from C to A in position 13,692 that leads to a proline (P) to glutamine (Q) change. Base A is present in 43.8% of reads while the reference C is present in 18.8%. The deamination C-to-T is present in three reads in terminal positions. In (b) a leucine (L) to phenylalanine (F) change (bases G to C) in position 13,858 in which two C-to-T transitions are present. In (c) a bona fide C-to-T change in which the frequency of T is above the expected by pure deamination level and therefore characterizing a proline (P) to leucine (L) change in position 13,704.
