## Supplementary material for "Reconstructing prehistoric viral genomes from Neanderthal sequencing data": Supplementary Figure S11.pdf

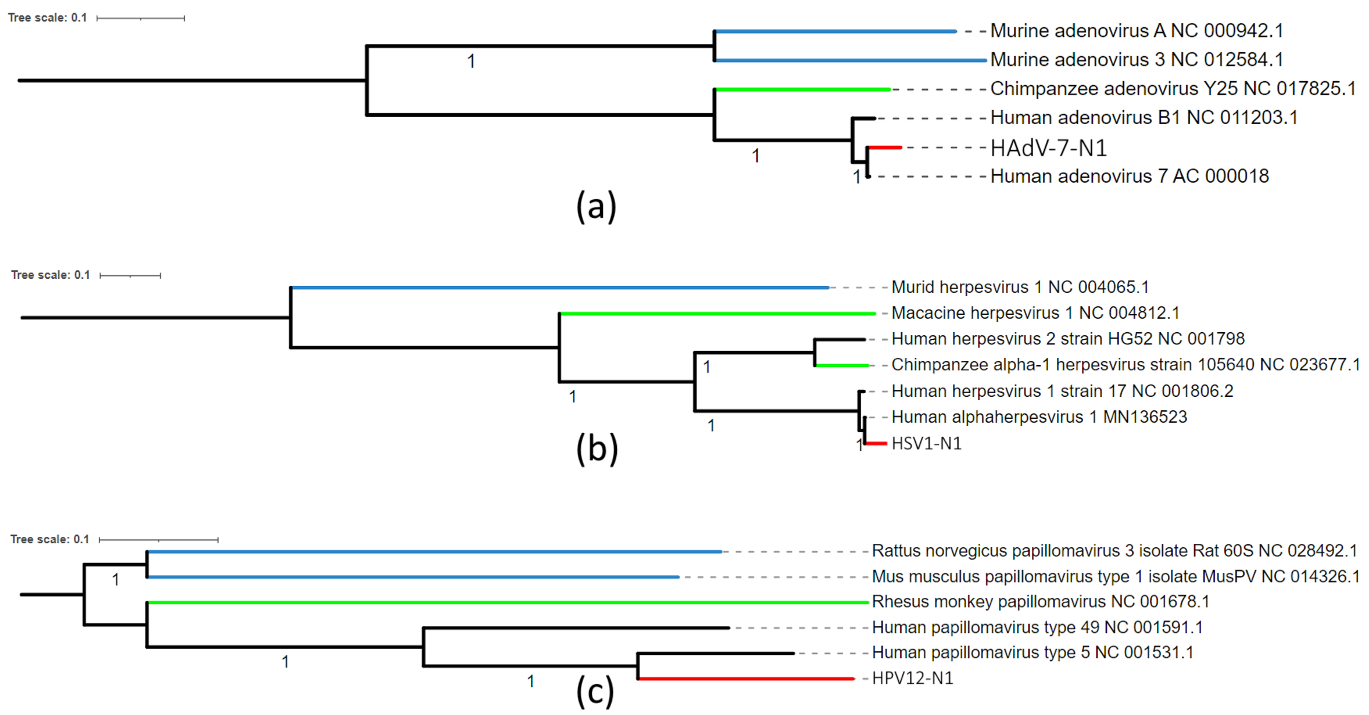


**Supplementary Figure S11.** Phylogenies of Neanderthal viruses HAdV7-N1, HSV1-N1 and HPV12-N1 compared to NCBI RefSeq primate and murid sequences. Red branches indicate Neanderthal inferred sequences, green branches indicate non-human primate sequences and blue branches indicate murid sequences. Trees were inferred from MAFFT alignments using a Maximum Likelihood model as implemented in FastTree 2.1.11 with GTR model, 4 categories of substitution rates and branch support by Shimodaira-Hasegawa test (Price et al., 2010, *PLoS ONE* 5(3): e9490. https://doi.org/10.1371/journal.pone.0009490). Tree scales (top left of each tree) indicate number of substitutions/sequence position.
