## Supplementary material for "Reconstructing prehistoric viral genomes from Neanderthal sequencing data": Supplementary Figure S12.pdf

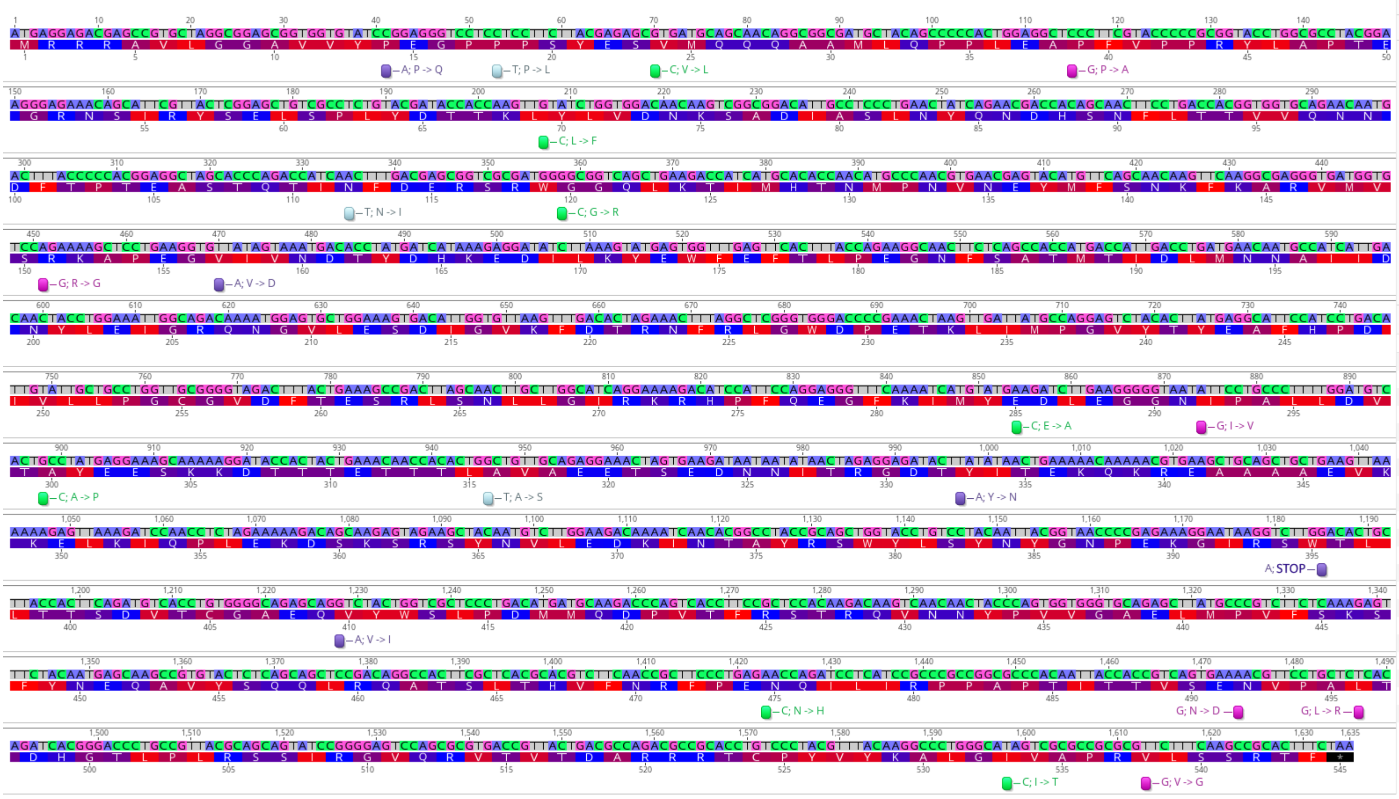


(a) L2

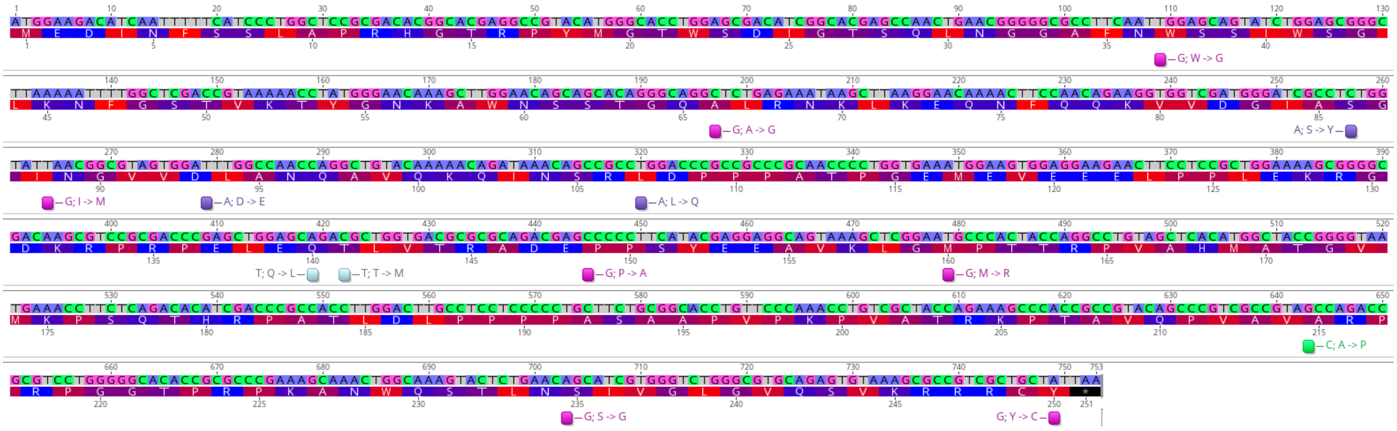


(b) L3

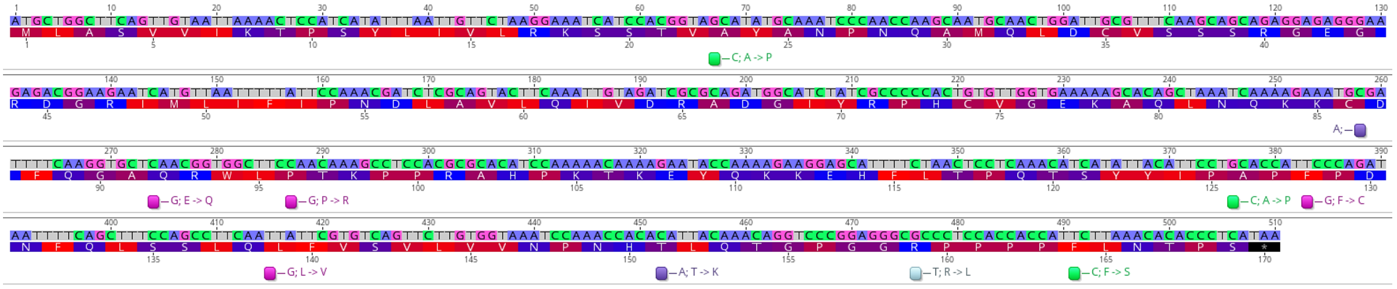


(c) L5

**Supplementary Figure S12.** Non-synonymous SNPs in adenovirus L2 gene (a), L3 gene (b) and L5 gene (c) of HAdV-7-N1 as compared to adenovirus assembly reference KX897164. Bases A=purple, C=green, G=pink, T-gray. Amino acid colors indicate red as the most hydrophobic (hydrophobicity=1), blue the most hydrophilic (hydrophobicity=0) and purple as intermediate (hydrophobicity≈0.5) (https://web.expasy.org/protscale/pscale/Hphob.Black.html). The original base is in the reference, the altered Neanderthal base is indicated by the color box and the amino acid change next to the changed base.
