## Supplementary material for "Reconstructing prehistoric viral genomes from Neanderthal sequencing data": Supplementary Figure S13.pdf

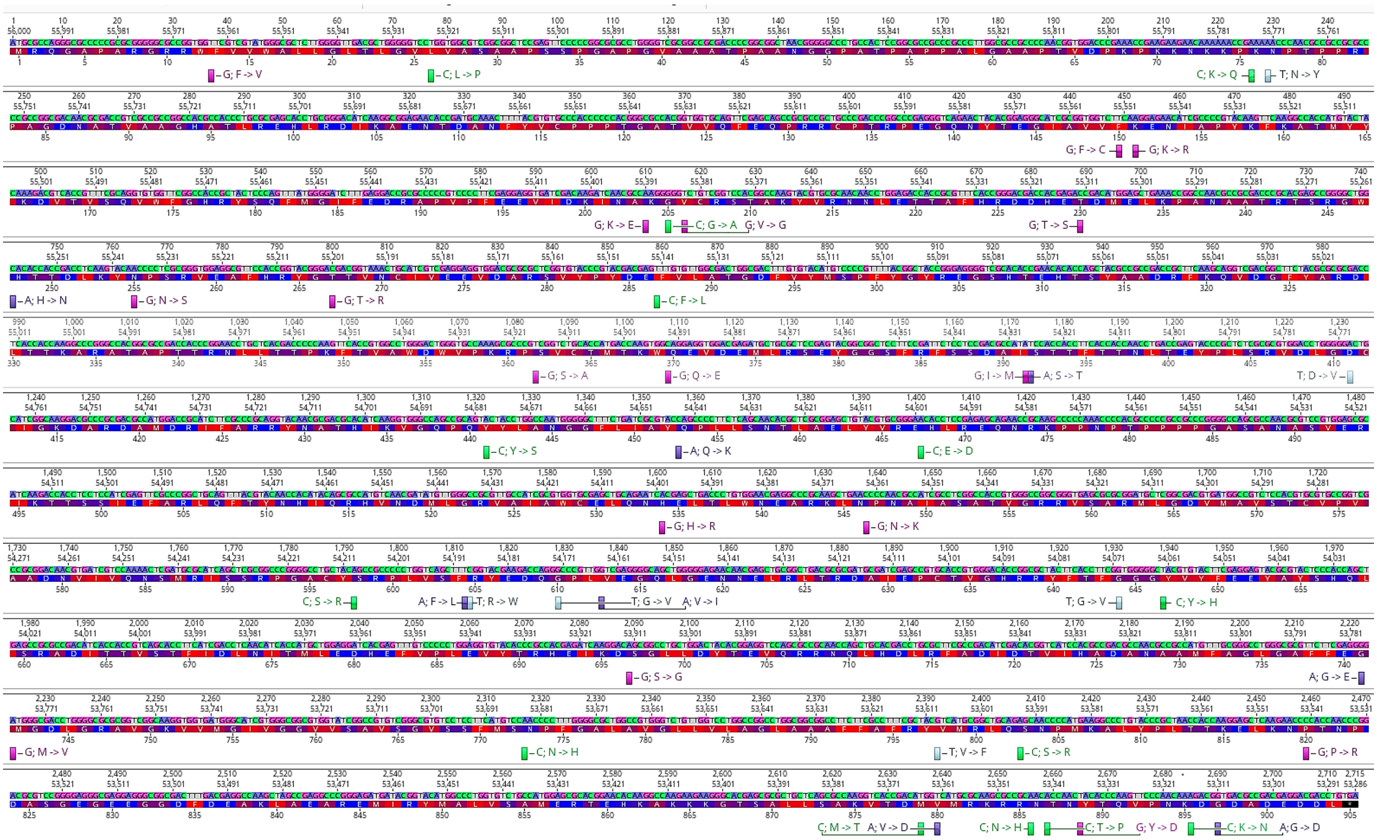

(a) UL27 - gB

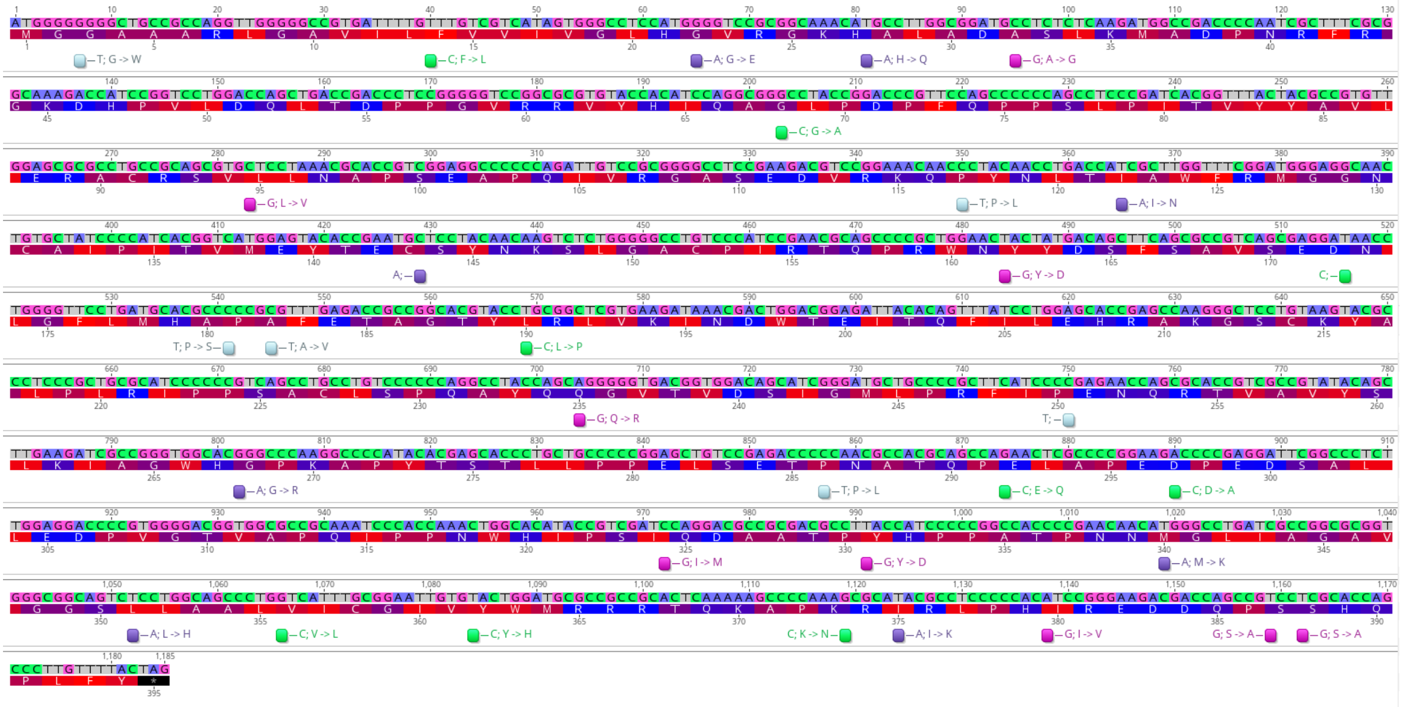

(b) US6 - gD

**Supplementary Figure S13.** Non-synonymous SNPs in herpesvirus UL27 gene (gB) (a) and US6 gene (gD) (b) of HSV1-N1 as compared to herpesvirus assembly reference MN136523. Bases A=purple, C=green, G=pink, T-gray. Amino acid colors indicate red as the most hydrophobic (hydrophobicity=1), blue the most hydrophilic (hydrophobicity=0) and purple as intermediate (hydrophobicity≈0.5) (https://web.expasy.org/protscale/pscale/Hphob.Black.html). The original base is in the reference, the altered Neanderthal base is indicated by the color box and the amino acid change next to the changed base.
