## Supplementary material for "Reconstructing prehistoric viral genomes from Neanderthal sequencing data": Supplementary Figure S14.pdf

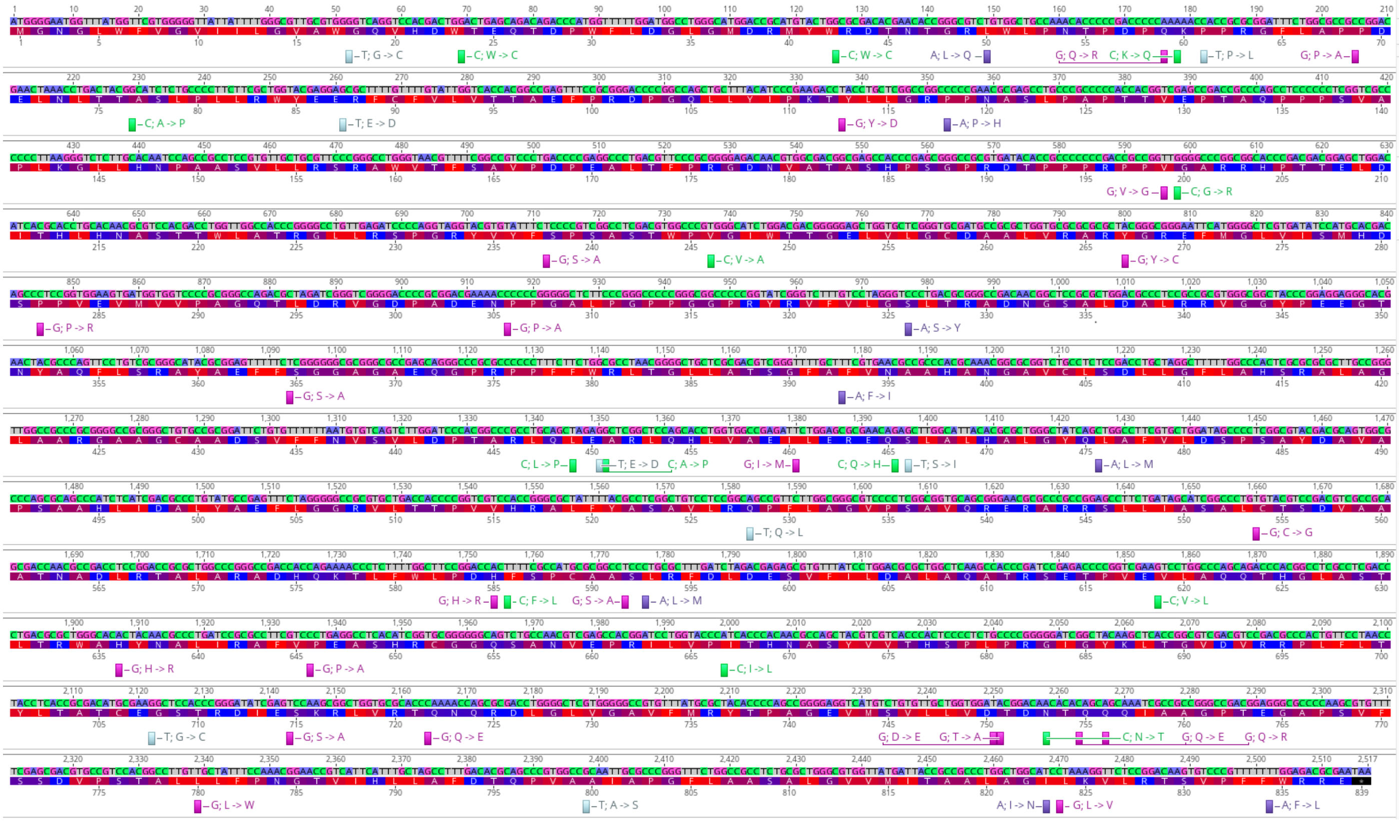

(a) UL22 – gH

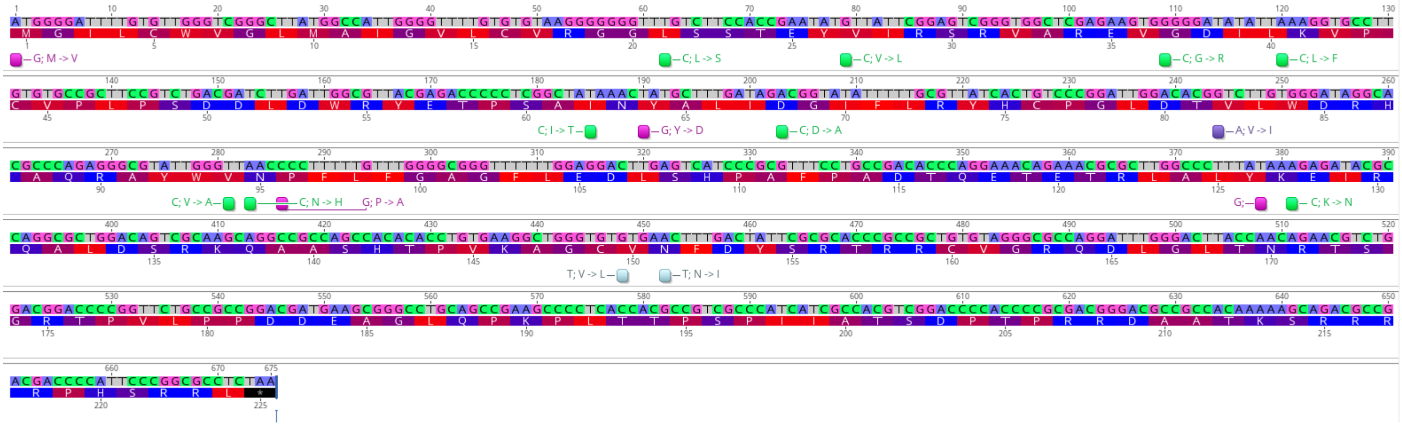

(b) UL1 - gL

**Supplementary Figure S14.** Non-synonymous SNPs in herpesvirus UL22 gene (gH) (a) and UL1 gene (gL) (b) of HSV1-N1 as compared to herpesvirus assembly reference MN136523. Bases A=purple, C=green, G=pink, T-gray. Amino acid colors indicate red as the most hydrophobic (hydrophobicity=1), blue the most hydrophilic (hydrophobicity=0) and purple as intermediate (hydrophobicity≈0.5) (https://web.expasy.org/protscale/pscale/Hphob.Black.html). The original base is in the reference, the altered Neanderthal base is indicated by the color box and the amino acid change next to the changed base.
