## Supplementary material for "Reconstructing prehistoric viral genomes from Neanderthal sequencing data": Supplementary Figure S15.pdf

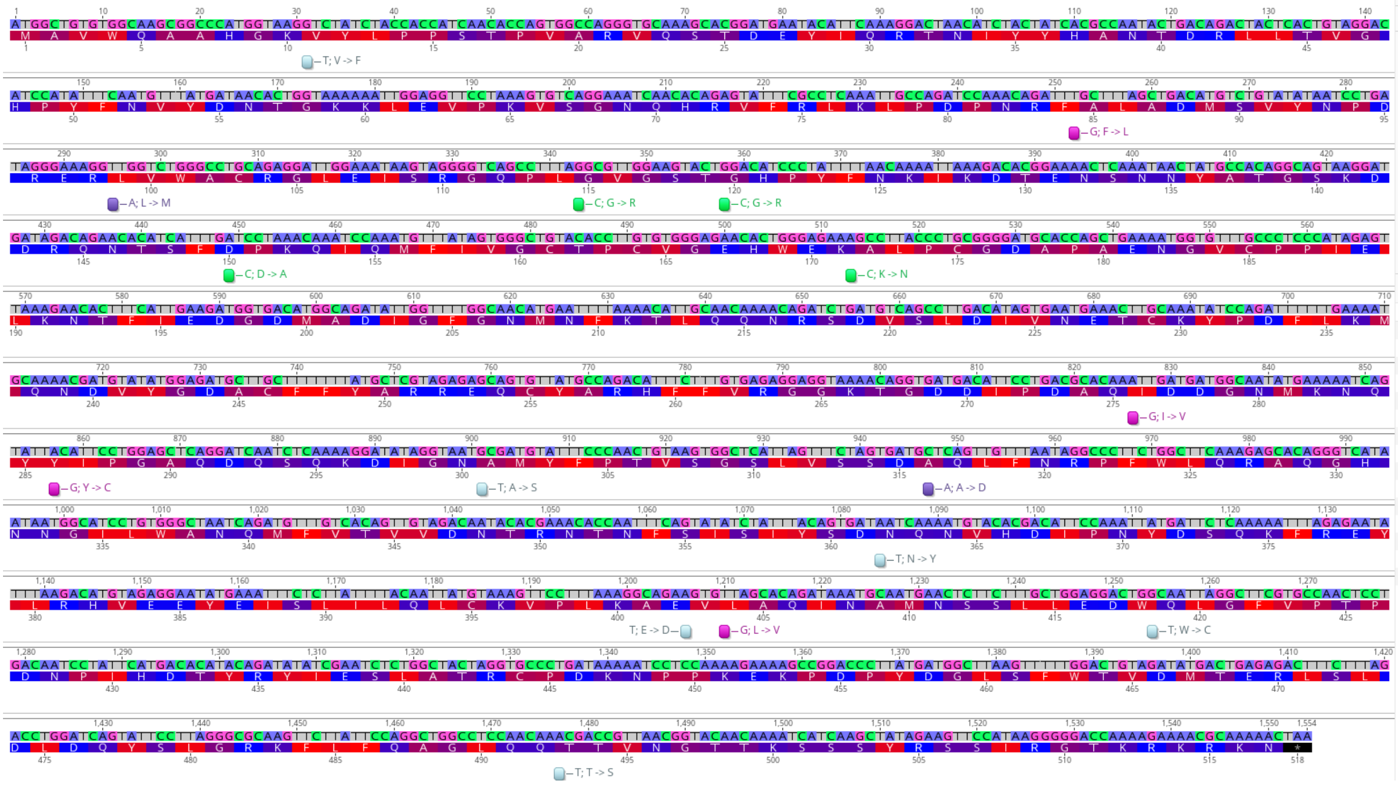

(a) L1

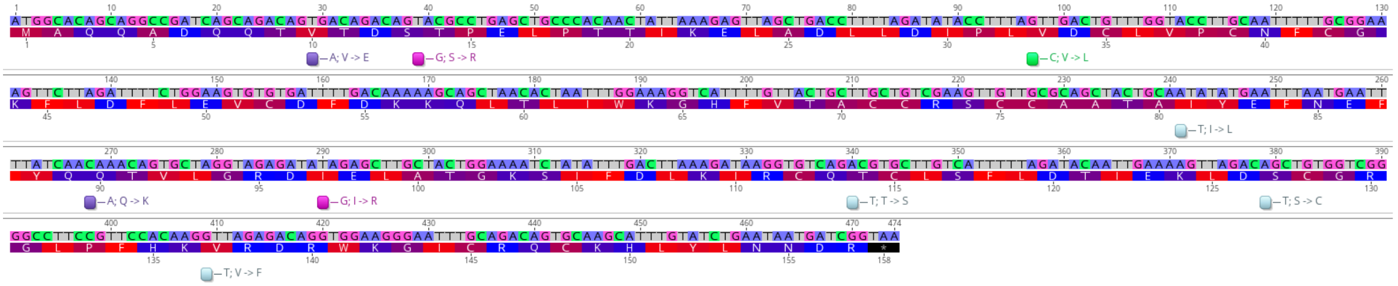

(b) E6

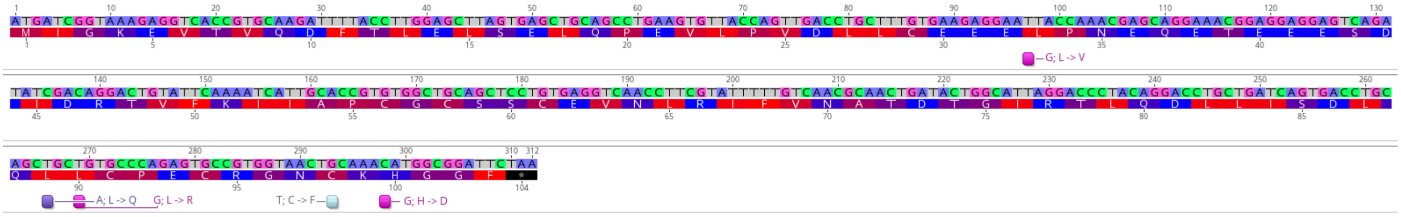

(c) E7

**Supplementary Figure S15.** Non-synonymous SNPs in papillomavirus L1 gene (a), E6 gene (b) and E7 gene (c) of HPV12-N1 as compared to papillomavirus assembly reference X74466. Bases A=purple, C=green, G=pink, T-gray. Amino acid colors indicate red as the most hydrophobic (hydrophobicity=1), blue the most hydrophilic (hydrophobicity=0) and purple as intermediate (hydrophobicity≈0.5) (https://web.expasy.org/protscale/pscale/Hphob.Black.html). The original base is in the reference, the altered Neanderthal base is indicated by the color box and the amino acid change next to the changed base.
