## Supplementary material for "Reconstructing prehistoric viral genomes from Neanderthal sequencing data": Supplementary Figure S16.pdf

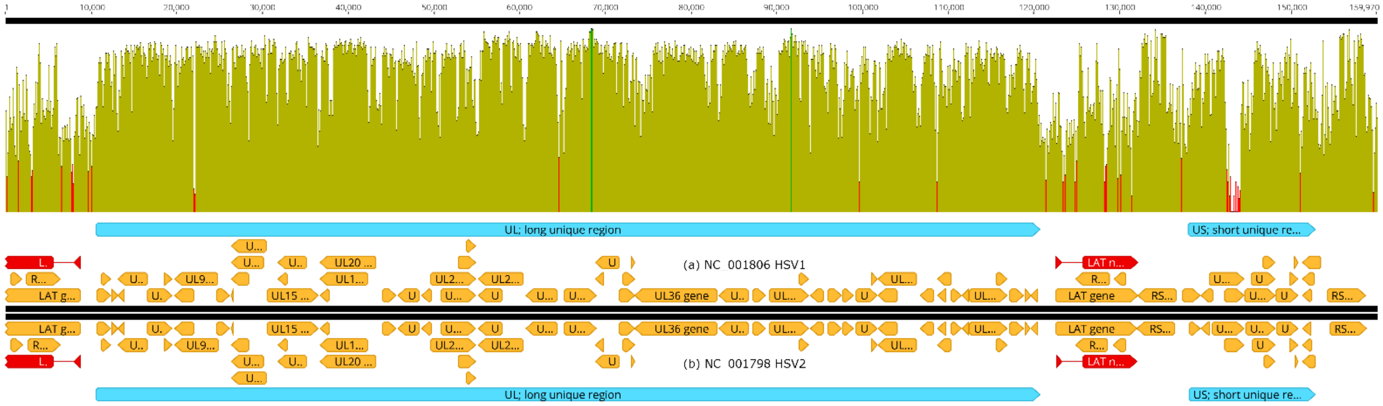


**Supplementary Figure S16.** Alignment of HSV1 and HSV2 reference genomes (NC_001806.2 and NC_001798.2). The global nucleotide similarity between these two viral genomes is 73%. A conserved region with 80% nucleotide identity, encompassing the long unique region (UL), is flanked by upstream variable regions with 50% nucleotide identity and downstream with 56% nucleotide identity. Dark green vertical bars indicate 100% similarity, olive green vertical bars indicate nucleotide identity <100% and ≥30%, red vertical bars indicate <30% identity. Orange arrows indicate genes, red arrows indicate noncoding RNAs, and light blue arrows indicate long (UL) and short (US) unique regions. Alignment with MAFFT (version 7.490) with scoring matrix PAM 200 / k=2, gap open penalty = 1.53 and offset value = 0.123 [35].
