## Supplementary material for "Reconstructing prehistoric viral genomes from Neanderthal sequencing data": Supplementary Table 1 Adenovirus Modeltest BIC.docx

Supplementary Table 1. Parameter estimates by ModelTest-NG and Bayesian Information Criterion for the adenovirus dataset.

| BIC | model | K | lnL | score | delta | weight |
| --- | --- | --- | --- | --- | --- | --- |
|  | 1 F81+I+G4 | 5 | -55676.6874 | 113485.8682 | 0.0000 | 0.8089 |
|  | 2 HKY+I+G4 | 6 | -55672.9996 | 113488.9460 | 3.0778 | 0.1736 |
|  | 3 TPM2uf+I+G4 | 7 | -55671.2055 | 113495.8112 | 9.9430 | 0.0056 |
|  | 4 TPM3uf+I+G4 | 7 | -55671.2152 | 113495.8306 | 9.9624 | 0.0056 |
|  | 5 TrN+I+G4 | 7 | -55671.3870 | 113496.1743 | 10.3061 | 0.0047 |
|  | 6 TPM1uf+I+G4 | 7 | -55672.8428 | 113499.0859 | 13.2177 | 0.0011 |
|  | 7 TIM2+I+G4 | 8 | -55669.2391 | 113502.3318 | 16.4636 | 0.0002 |
|  | 8 TIM3+I+G4 | 8 | -55669.5951 | 113503.0439 | 17.1757 | 0.0002 |
|  | 9 JC+I+G4 | 2 | -55701.2196 | 113503.5724 | 17.7042 | 0.0001 |
|  | 10 K80+I+G4 | 3 | -55696.8948 | 113505.3763 | 19.5081 | 0.0000 |

Best model according to BIC

| Model: | F81+I+G4 |  |
| --- | --- | --- |
| lnL: | -55676.6874 |  |
| Frequencies: | 0.2515 0.2594 | 0.2542 0.2348 |
| Subst. Rates: | 1.0000 1.0000 | 1.0000 1.0000 1.0000 1.0000 |
| Inv. sites prop: | 0.5040 |  |
| Gamma shape: | 0.1924 |  |
| Score: | 113485.8682 |  |
| Weight: | 0.8089 |  |

Parameter importances P.Inv:	-

```
Gamma:	-
Gamma-Inv:	1.0000
Frequencies:	0.9998
```

Model averaged estimates P.Inv:	-

```
Alpha:	-
Alpha-P.Inv:	0.2345
P.Inv-Alpha:	0.5312
Frequencies:	0.2515 0.2594 0.2542 0.2348
```
