## Supplementary material for "Reconstructing prehistoric viral genomes from Neanderthal sequencing data": Supplementary Table 2 Herpesvirus Modeltest BIC.docx

Supplementary Table 2. Parameter estimates by ModelTest-NG and Bayesian Information Criterion for the herpesvirus dataset.

| BIC | model | K | lnL | score | delta | weight |
| --- | --- | --- | --- | --- | --- | --- |
|  | 1 GTR+I+G4 | 10 | -370860.6877 | 744165.4747 | 0.0000 | 0.9905 |
|  | 2 TVM+I+G4 | 9 | -370871.2967 | 744174.7703 | 9.2956 | 0.0095 |
|  | 3 TIM1+I+G4 | 8 | -370961.0703 | 744342.3950 | 176.9202 | 0.0000 |
|  | 4 TPM1uf+I+G4 | 7 | -370970.8451 | 744350.0223 | 184.5475 | 0.0000 |
|  | 5 TIM3+I+G4 | 8 | -371646.0510 | 745712.3565 | 1546.8817 | 0.0000 |
|  | 6 TPM3uf+I+G4 | 7 | -371659.9747 | 745728.2815 | 1562.8067 | 0.0000 |
|  | 7 TIM2+I+G4 | 8 | -371764.7860 | 745949.8265 | 1784.3518 | 0.0000 |
|  | 8 TPM2uf+I+G4 | 7 | -371772.1024 | 745952.5367 | 1787.0620 | 0.0000 |
|  | 9 TrN+I+G4 | 7 | -371985.5530 | 746379.4379 | 2213.9632 | 0.0000 |
|  | 10 HKY+I+G4 | 6 | -371994.2478 | 746384.9051 | 2219.4304 | 0.0000 |

Best model according to BIC

| Model: | GTR+I+G4 |
| --- | --- |
| lnL: | -370860.6877 |
| Frequencies: | 0.1599 0.3371 0.3438 0.1592 |
| Subst. Rates: | 0.9903 1.8152 0.6142 0.3307 2.0155 1.0000 |
| Inv. sites prop: | 0.8408 |
| Gamma shape: | 0.2714 |
| Score: | 744165.4747 |
| Weight: | 0.9905 |

Parameter importances P.Inv:	-

```
Gamma:	-
Gamma-Inv:	1.0000
Frequencies:	1.0000
```

Model averaged estimates P.Inv:	-

```
Alpha:	-
Alpha-P.Inv:	0.2714
P.Inv-Alpha:	0.8408
Frequencies:	0.1599 0.3371 0.3438 0.1592
```
