## Supplementary material for "Reconstructing prehistoric viral genomes from Neanderthal sequencing data": Supplementary Table 3 Papillomavirus Modeltest BIC.docx

Supplementary Table 3. Parameter estimates by ModelTest-NG and Bayesian Information Criterion for the papillomavirus dataset.

```
BIC       model              K            lnL          score          delta    weight
--------------------------------------------------------------------------------
       1  GTR+I+G4          10    -58362.0041    118089.9482         0.0000    1.0000
       2  TIM2+I+G4          8    -58383.9226    118115.9297        25.9815    0.0000
       3  TVM+I+G4           9    -58379.6185    118116.2493        26.3011    0.0000
       4  TPM2uf+I+G4        7    -58397.0877    118133.3322        43.3840    0.0000
       5  TIM3+I+G4          8    -58393.3002    118134.6849        44.7367    0.0000
       6  TPM3uf+I+G4        7    -58406.7987    118152.7542        62.8060    0.0000
       7  TrN+I+G4           7    -58410.1869    118159.5307        69.5825    0.0000
       8  TIM1+I+G4          8    -58406.0633    118160.2111        70.2629    0.0000
       9  HKY+I+G4           6    -58419.6900    118169.6090        79.6609    0.0000
      10  TPM1uf+I+G4        7    -58415.6606    118170.4781        80.5299    0.0000
--------------------------------------------------------------------------------
Best model according to BIC
---------------------------
Model:              GTR+I+G4
lnL:                -58362.0041
Frequencies:        0.3216 0.1878 0.2134 0.2772
Subst. Rates:       1.9422 3.8666 1.6240 1.6263 5.2034 1.0000
Inv. sites prop:    0.3572
Gamma shape:        1.0434
Score:              118089.9482
Weight:             1.0000
---------------------------
Parameter importances
---------------------------
P.Inv:              -
Gamma:              -
Gamma-Inv:          1.0000
Frequencies:        1.0000
---------------------------
Model averaged estimates
---------------------------
P.Inv:              -
Alpha:              -
Alpha-P.Inv:        1.0434
P.Inv-Alpha:        0.3572
Frequencies:        0.3216 0.1878 0.2134 0.2772
```
