## Supplementary material for "Reconstructing prehistoric viral genomes from Neanderthal sequencing data": Supplementary Table 10.pdf

**Supplementary Table 10**. Similarity matrix of human, primate, and murid adenovirus NCBI RefSeq sequences.

| Adenovirus sequences | Human adenovirus 7 AC_000018 | HAdV-7-N1 consensus | Human adenovirus B1 NC_011203.1 | Chimpanzee adenovirus Y25 NC_017825.1 | Murine adenovirus 3 NC_012584.1 | Murine adenovirus A NC_000942.1 |
|---|---|---|---|---|---|---|
| Human adenovirus 7 AC_000018 | 100.0 | 94.7 | 95.3 | 71.3 | 40.4 | 41.3 |
| HAdV-7-N1 Consensus | 94.7 | 100.0 | 92.0 | 69.5 | 40.0 | 40.8 |
| Human adenovirus B1 NC_011203.1 | 95.3 | 92.0 | 100.0 | 71.6 | 40.4 | 41.4 |
| Chimpanzee adenovirus Y25 NC_017825.1 | 71.3 | 69.5 | 71.6 | 100.0 | 39.0 | 40.0 |
| Murine adenovirus 3 NC_012584.1 | 40.4 | 40.0 | 40.4 | 39.0 | 100.0 | 64.4 |
| Murine adenovirus A NC_000942.1 | 41.3 | 40.8 | 41.4 | 40.0 | 64.4 | 100.0 |
