## Supplementary material for "Reconstructing prehistoric viral genomes from Neanderthal sequencing data": Supplementary Table 11.pdf

**Supplementary Table 11**. Identity (%) matrix of human, primate, and murid herpesvirus NCBI RefSeq sequences.

| Herpesvirus sequences | HSV1-N1 consensus | Human alphaherpesvirus 1 MN136523 | Human herpesvirus 1 strain 17 NC_001806.2 | Chimpanzee alpha-1 herpesvirus strain 105640 NC_023677.1 | Human herpesvirus 2 strain HG52 NC_001798 | Macacine herpesvirus 1 NC_004812.1 | Murid herpesvirus 1 NC_004065.1 |
|---|---|---|---|---|---|---|---|
| HSV1-N1 consensus | 100.0 | 95.7 | 94.5 | 70.5 | 70.1 | 56.6 | 28.6 |
| Human alphaherpesvirus 1 MN136523 | 95.7 | 100.0 | 97.7 | 72.1 | 71.8 | 57.9 | 29.0 |
| Human herpesvirus 1 strain 17 NC_001806.2 | 94.5 | 97.7 | 100.0 | 72.3 | 72.1 | 57.8 | 29.0 |
| Chimpanzee alpha-1 herpesvirus strain 105640 NC_023677.1 | 70.5 | 72.1 | 72.3 | 100.0 | 88.4 | 58.2 | 29.2 |
| Human herpesvirus 2 strain HG52 NC_001798 | 70.1 | 71.8 | 72.1 | 88.4 | 100.0 | 59.0 | 29.5 |
| Macacine herpesvirus 1 NC_004812.1 | 56.6 | 57.9 | 57.8 | 58.2 | 59.0 | 100.0 | 29.6 |
| Murid herpesvirus 1 NC_004065.1 | 28.6 | 29.0 | 29.0 | 29.2 | 29.5 | 29.6 | 100.0 |
