## Supplementary material for "Reconstructing prehistoric viral genomes from Neanderthal sequencing data": Supplementary Table 12.pdf

**Supplementary Table 12.** Identity (%) matrix of human, primate, and murid papillomavirus NCBI RefSeq sequences.

| Papillomavirus Sequences | HPV12-N1 consensus | Human papillomavirus type 5 NC_001531.1 | Human papillomavirus type 49 NC_001591.1 | *Mus musculus* papillomavirus type 1 isolate MusPV NC_014326.1 | *Rattus norvegicus* papillomavirus 3 isolate Rat_60S NC_028492.1 | *Rhesus* monkey papillomavirus NC_001678.1 |
|---|---|---|---|---|---|---|
| HPV12-N1 consensus | 100.0 | 74.0 | 59.3 | 41.9 | 42.9 | 39.7 |
| Human papillomavirus type 5 NC_001531.1 | 74.0 | 100.0 | 61.1 | 42.4 | 44.0 | 40.1 |
| Human papillomavirus type 49 NC_001591.1 | 59.3 | 61.1 | 100.0 | 43.2 | 44.5 | 41.1 |
| *Mus musculus* papillomavirus type 1 isolate MusPV NC_014326.1 | 41.9 | 42.4 | 43.2 | 100.0 | 43.7 | 39.4 |
| *Rattus norvegicus* papillomavirus 3 isolate Rat_60S NC_028492.1 | 42.9 | 44.0 | 44.5 | 43.7 | 100.0 | 39.7 |
| *Rhesus* monkey papillomavirus NC_001678.1 | 39.7 | 40.1 | 41.1 | 39.4 | 39.7 | 100.0 |
