## Supplementary figures and images for "Reconstructing prehistoric viral genomes from Neanderthal sequencing data"

### Supplementary Data S16 - Krona view of ERR10073060 Run.png

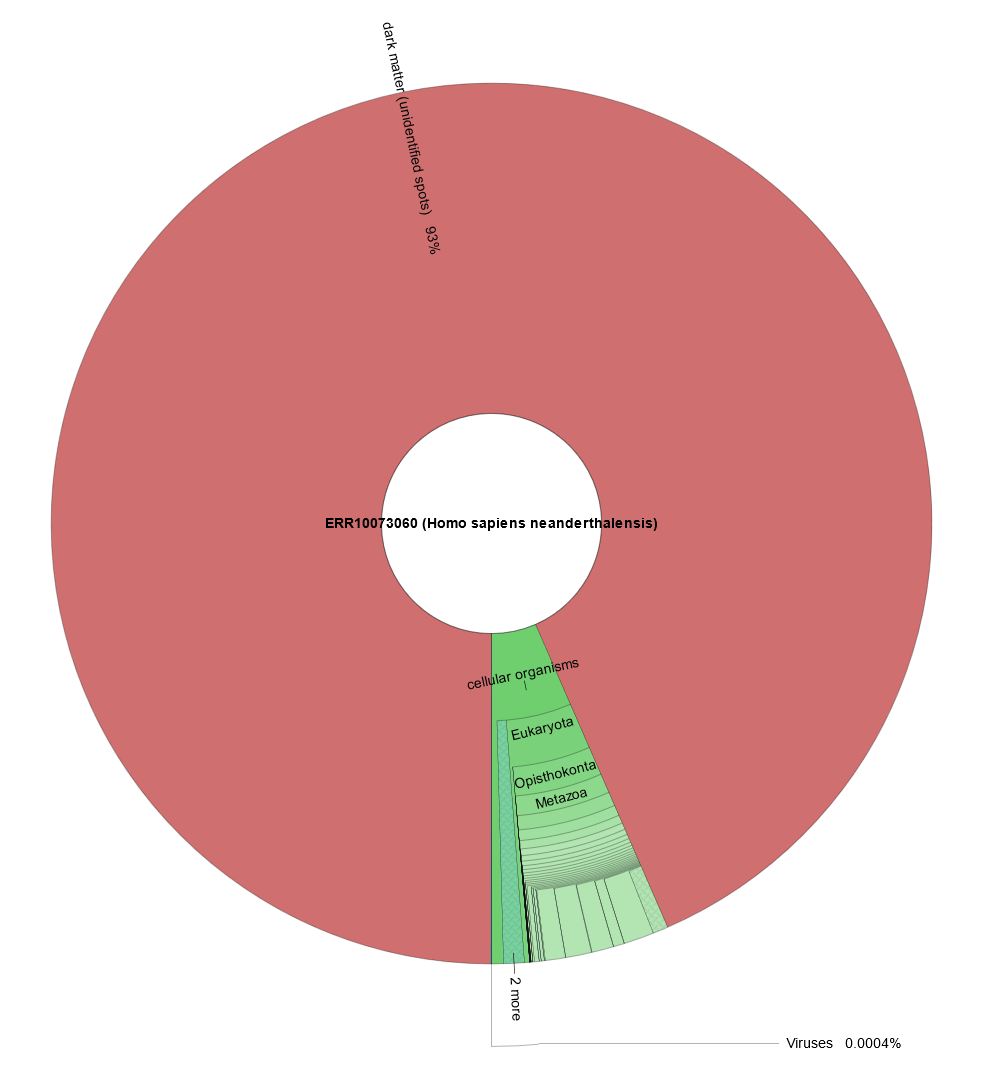

### Supplementary Data S17 - ERR10073060 Run Browser SRA Archive NCBI.png

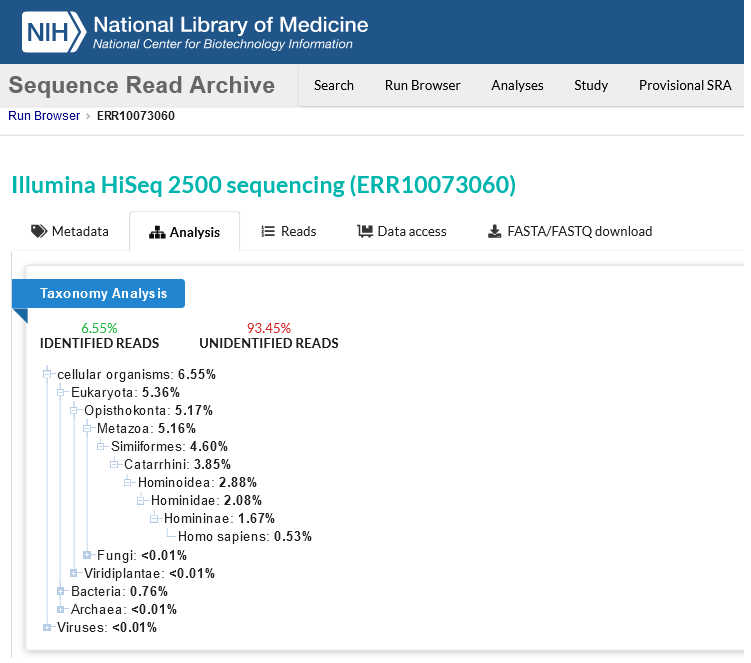

### Supplementary Figure S1 Adenovirus NJ-MegaBlast.png

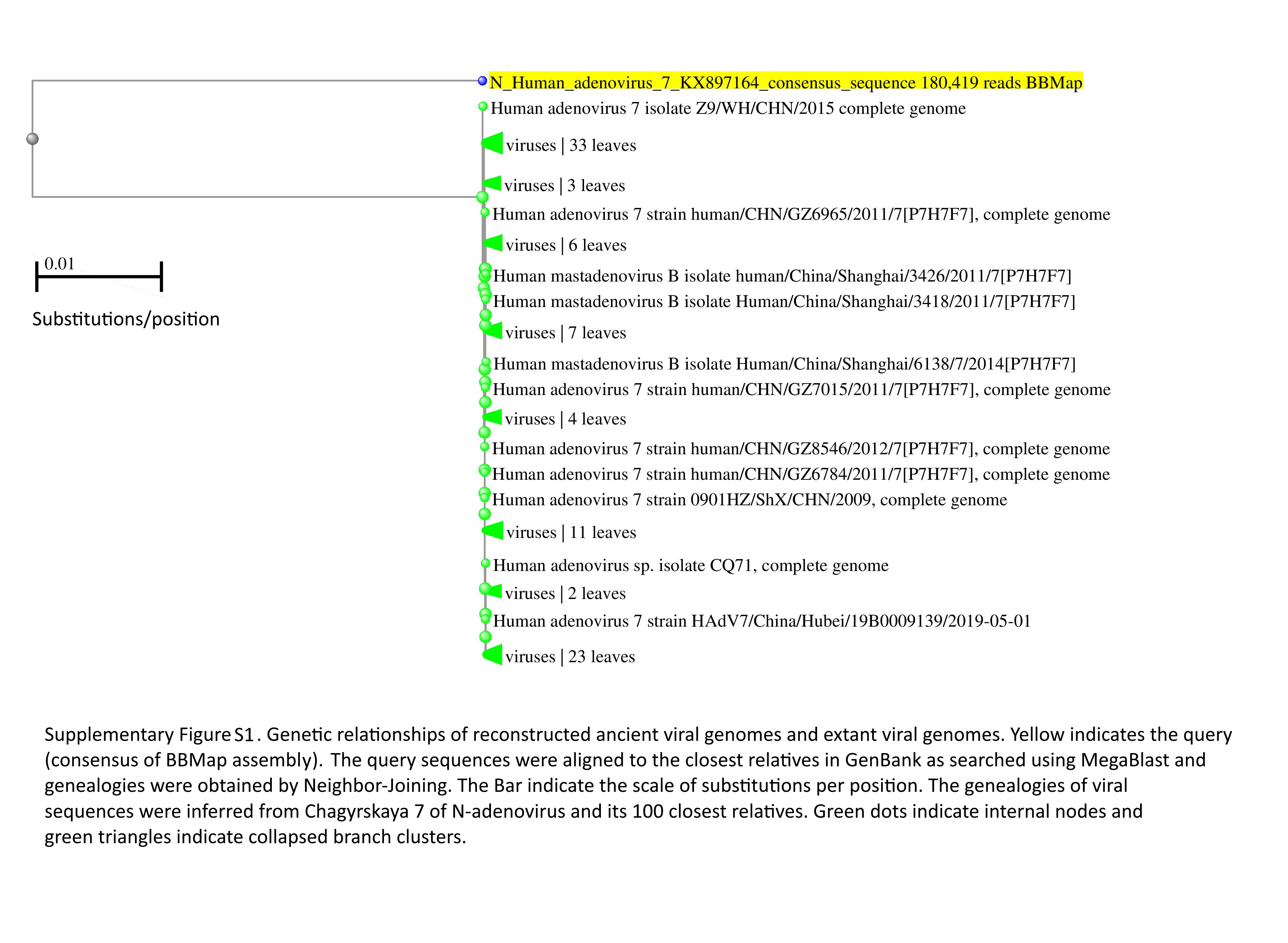

### Supplementary Figure S2 Herpesvirus NJ-MegaBlast.png

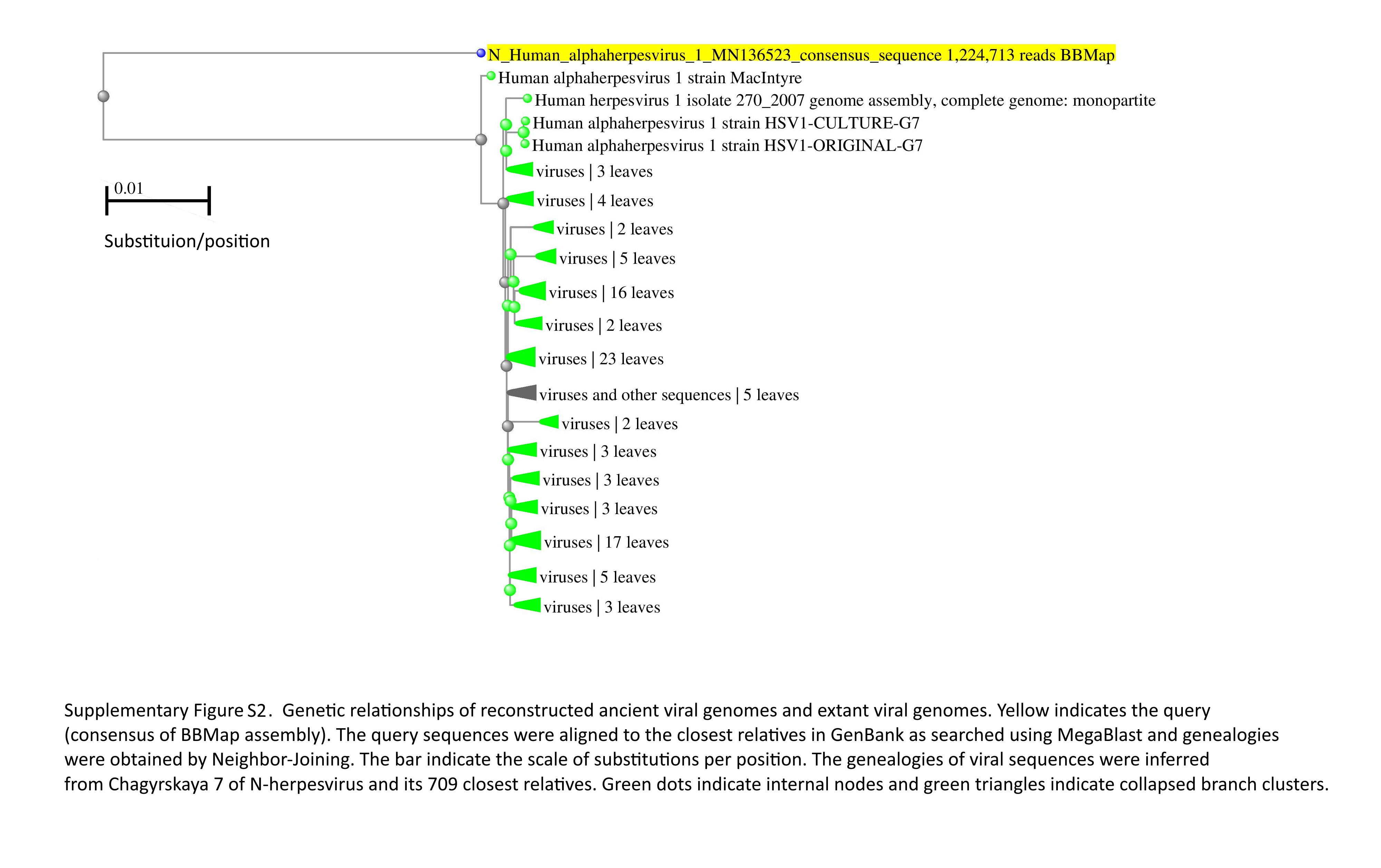

### Supplementary Figure S3 Papillomavirus NJ-MegaBlast.png

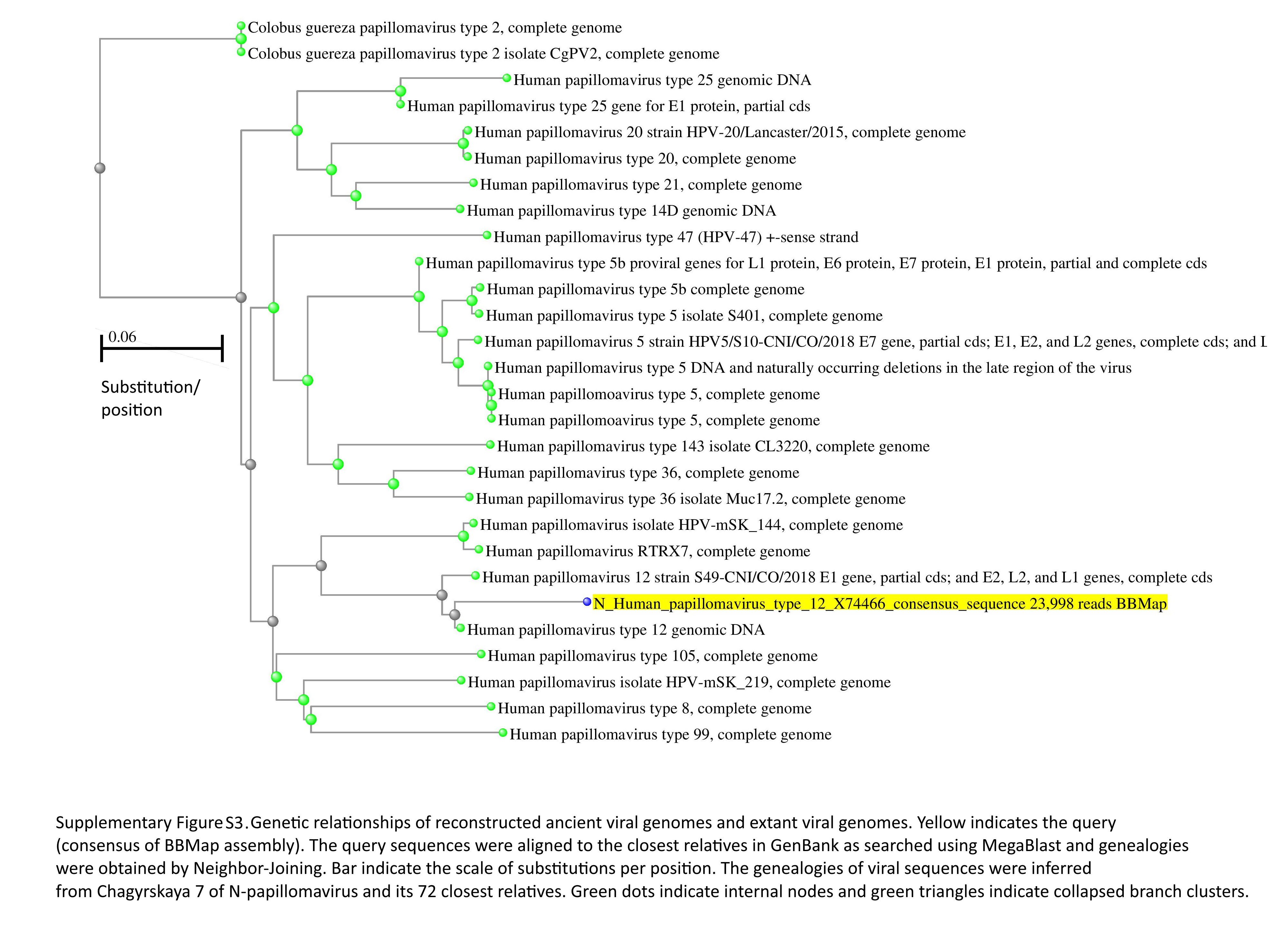

### Supplementary Figure S4 Adenovirus ML Tree.jpg

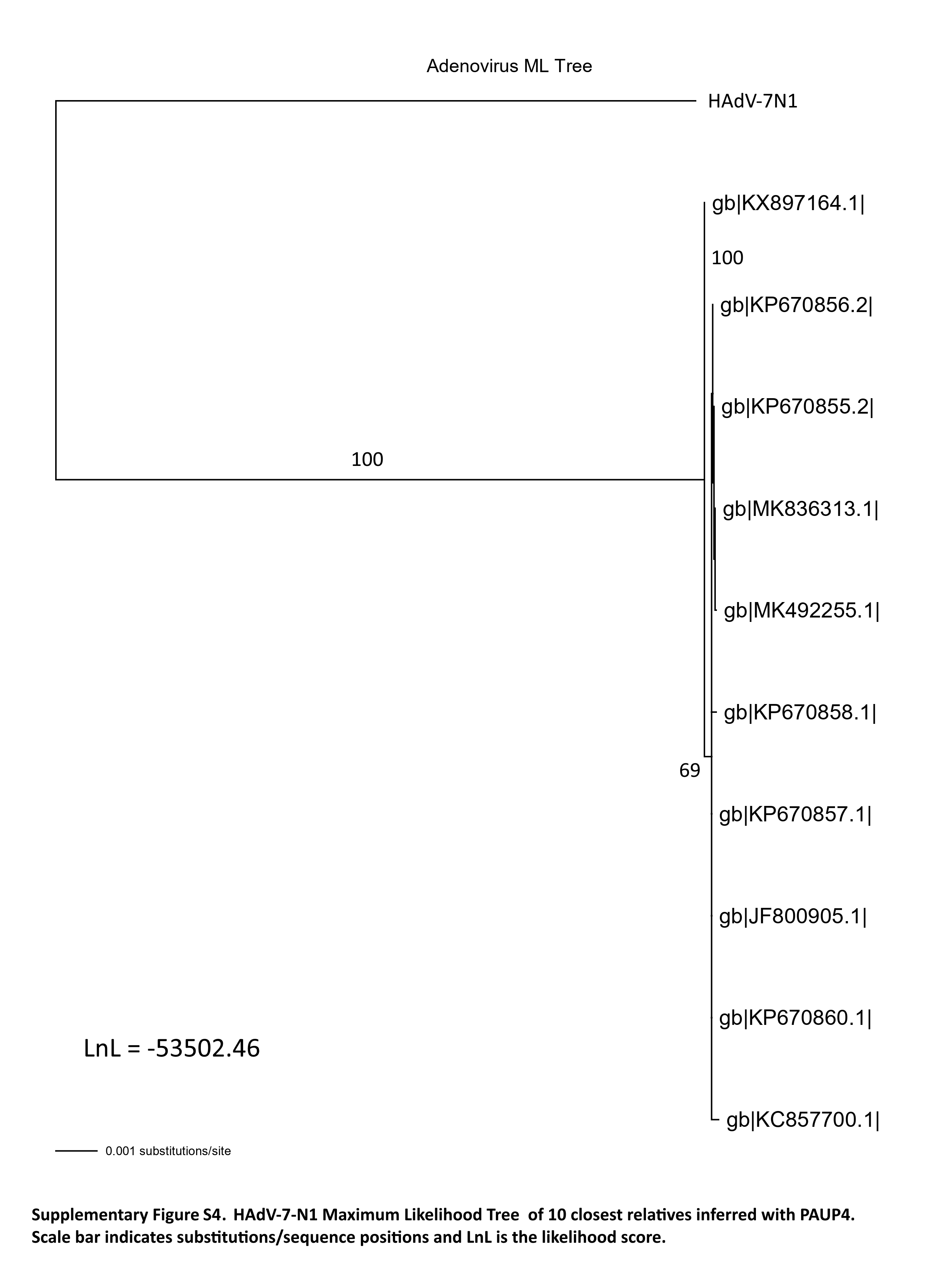

### Supplementary Figure S5 Herpesvirus ML Tree.jpg

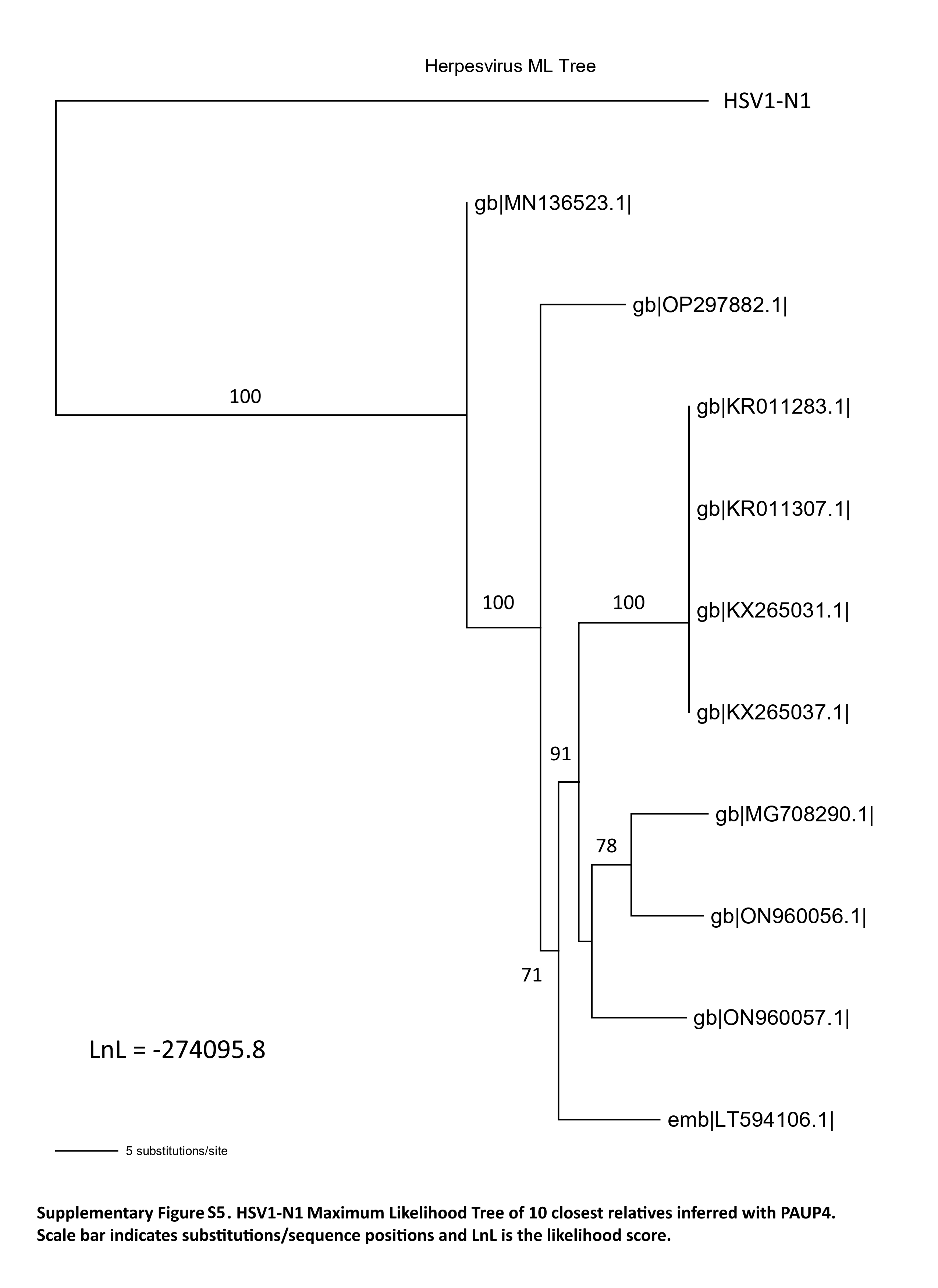

### Supplementary Figure S6 Papillomavirus ML Tree.jpg

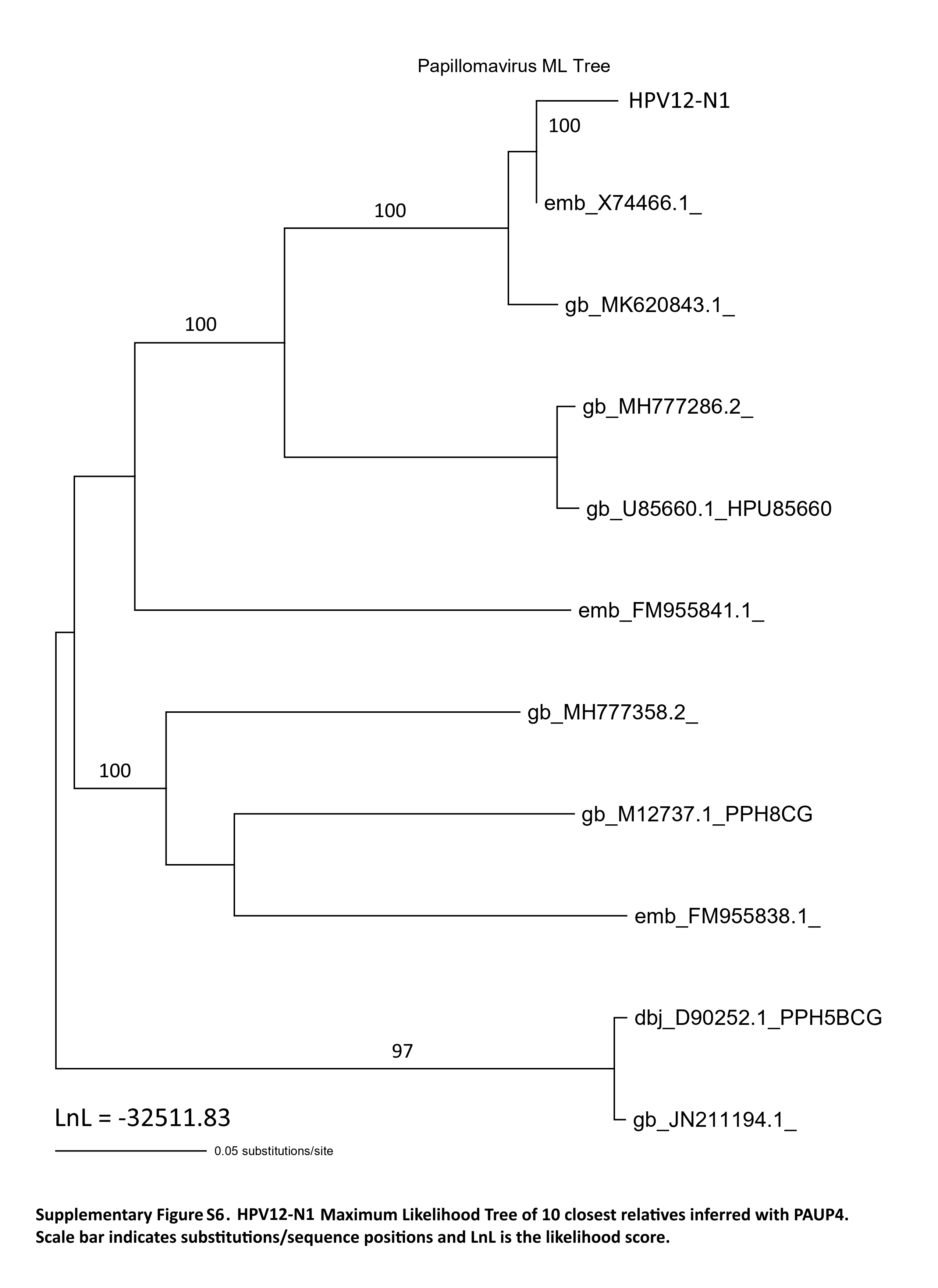
